# DiffDec: Structure-Aware Scaffold Decoration with an End-to-End Diffusion Model

**DOI:** 10.1101/2023.10.08.561377

**Authors:** Junjie Xie, Sheng Chen, Jinping Lei, Yuedong Yang

## Abstract

In molecular optimization, one popular way is R-groups decoration on molecular scaffolds, and many efforts have been put to generate R-groups based on deep generative models. However, these methods mostly use information of known binding ligands, without fully utilizing target structure information. In this study, we proposed a new method, DiffDec, to involve three-dimensional pocket constraints by a modified diffusion technique for optimizing molecules through molecular scaffold decoration. For an end-to-end generation of R-groups with different sizes, we designed a novel fake atom mechanism. DiffDec was shown able to generate structure-aware R-groups, and simultaneously generate multiple R-groups for one scaffold on different growth anchors. The growth anchors could be provided by users or automatically determined by our model. DiffDec achieved R-group recovery rates of 69.67% and 45.34% in the single and multiple R-group decoration tasks, respectively, and these values were significantly higher than competing methods (37.33% and 26.85%). According to the molecular docking study, our decorated molecules obtained better average binding affinity than baseline methods. The docking pose analysis revealed that DiffDec could decorate scaffolds with R-groups that exhibited improved binding affinities and more favourable interactions with the pocket. These results demonstrated the potential and applicability of DiffDec in real-world scaffold decoration for molecular optimization.

## Introduction

Computational methods have sped up the process of drug development in a cost-effective manner and minimized the risk of failure in the later stages.^1^ Existing methods can utilize known compounds to design novel compounds, also known as ligand-based drug design (LBDD).^2^ Traditional LBDD methods like pharmacophore model ^3^ or quantitative structure activity relationship model^4^ learn structure–activity relationship of ligands from bioactive data to design new drug candidates. ^5^ To efficiently explore the vast chemical space, deep generative models have gradually become the main engine in the majority of drug design tools.^6,7^ Based on different molecular representations, the existing molecular generative models can be roughly split into two categories: SMILES-based and graph-based. SMILES-based methods^8–12^ generally generate desired compounds through language models like long short-term memory^13^ and transformer,^14^ which usually have simple network architectures and can be trained fast. Differently, graph-based methods represent a molecule as a graph where its node-edge structure naturally matches the atom-bond information of molecules.^15–17^ Such methods have rapidly developed especially with the recently developed E(3)-equivariant graph neural network architectures^18^ that could embed the 3D molecular information.

While most of these deep generative models focus on *de novo* molecular generation, an important stage of drug design is lead optimization, which aims to optimize the favorable properties of compounds by changing part of their atoms.^19^ In particular, one important type of molecular optimization is scaffold decoration^20,21^ that retains the core scaffold and modifies molecules in specific motifs such as R-groups. Recently, LibINVENT^22^ leveraged chemical reaction templates to slice the molecules into scaffolds and decorations, and then trained a deep generative model to generate R-groups that decorated the extracted molecular scaffolds. This reaction-based R-group decoration method guarantees that the generated molecules are highly synthetic and accessible. However, this ligand-based generative model was based on recurrent neural networks and SMILES molecular representation, ^23^ which were not capable to take the 3D structure information of protein pockets into consideration.

In parallel to LBDD, structure-based drug design (SBDD)^24^ directly learns protein structure information, aiming to design drugs that selectively and specifically target disease-causing receptors. ^25^ Nowadays, there is a rapid increase in the accessible structural data because of the breakthrough in structural biology (e.g. X-ray crystallography^26^ and Cryo-EM^27^) and protein structure prediction (e.g. AlphaFold^28^ and ESMFold^29^), which empowers SBDD to be applied on most diseases with known protein targets. Traditional SBDD methods like molecular dynamics simulation^30^ and molecular docking^31^ are mostly used for virtual screening of known compounds but not good at designing novel molecules. Recently, increasing works attempted to incorporate 3D protein structures in molecule design tasks. For example, Pocket2Mol^32^ utilized equivariant graph neural networks to learn the chemical and geometric constraints imposed by protein pockets, and sampled molecules in an auto-regressive manner. FLAG^33^ generated 3D molecules using realistic substructures fragment-by-fragment around the binding pocket. Unfortunately, these auto-regressive generation methods are prone to suffering from inaccurate accumulation for unrealistic 3D molecular structures.^34^ As a full-atom generation manner rather than an auto-regressive manner, the diffusion model^35^ is recently applied to the field of small molecule generation. DiffLinker^36^ employed a conditional diffusion model and a linker size predictor to design molecular linkers for a set of fragments and proteins. DiffSBDD^37^ proposed an E(3)-equivariant 3D-conditional diffusion model to generate ligands conditioned on the binding pockets. However, such methods haven’t been used to decorate molecular scaffolds for optimization.

In this study, we have developed a new method, DiffDec, to optimize molecules through molecular scaffold decoration conditioned on the 3D protein pocket by an E(3)-equivariant graph neural network and diffusion model. Specifically, we constructed two 3D scaffold decoration datasets through reaction-based slicing of 3D protein-ligand complex in CrossDocked dataset.^38^ According to the scaffold and the context pocket, we trained a 3D conditional diffusion model to generate structure-aware R-groups. For an end-to-end generation of different-sized R-groups, we newly introduced a fake atom mechanism. Through comprehensive evaluations, DiffDec exhibited superior performance compared to baseline methods in terms of validity, R-group recovery, similarity, and binding affinity on the two benchmark datasets. Notably, DiffDec could identify the growth anchors and generate R-groups well for the scaffolds without provided anchors. More importantly, case studies revealed that the compounds decorated using DiffDec exhibited realistic 3D structures and more protein-ligand interactions compared to the reference molecules.

## Materials and methods

### Datasets

In the scaffold decoration dataset constructed by Fialková et al.,^22^ the sliced scaffolds and R-groups are represented by SMILES and the 3D structural information is unavailable. Therefore, we curated two benchmark datasets for single and multi R-groups scaffold decoration based on three-dimensional protein-ligand complex structures by using the Cross-Docked dataset^38^ and reaction-based slicing method. Specifically, the CrossDocked dataset included 22.5 million protein-ligand complexes at various levels of quality. Following the same data preparation and splitting strategies as in previous methods,^39^ we used 100,000 protein-ligand pairs for subsequent processing as the training set and 100 complexes from the remaining clusters as the test set. The split was done by 30% protein sequence identity using mmseqs2.^40^ The ligands were sliced into scaffolds and R-groups utilizing 37 customized reaction-based rules^22^ and ultimately the final training set and test set were obtained. This reaction-based division method improved the validity and synthetic accessibility of scaffold decoration with simple chemical reactions. For single R-group scaffold decoration dataset, we set the maximum cut in the configuration file of slicing molecules to 1. It contained 76,108 tuples with molecular scaffolds, pockets, and R-groups in training set and 43 tuples in test set. For a more challenging multi R-groups dataset, we changed the maximum cut to 4. It contained 154,792 tuples in training set and 102 tuples in test set. The detailed information of these two datasets is shown in Table S1 and Figure S1.

### Overview of DiffDec

DiffDec is capable of performing scaffold decoration under the context of pockets in 3D space. We use a set of atoms to represent molecules and proteins. Following the framework of denoising diffusion probabilistic model (DDPM), ^41^ atoms in the target pocket ***p*** and given scaffold ***s*** are represented as context 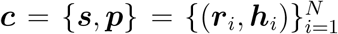, where ***r***_*i*_ is the 3D geometric coordinate of the *i*th heavy atom, ***h***_*i*_ is the corresponding element types and *N* is the number of atoms of the scaffold ***s*** and protein ***p***. As shown in Figure 1, our objective is to generate the decorations, i.e. R-groups 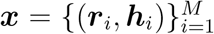 to form a complete molecule conditioned on the scaffold and protein pocket. The method consists of a diffusion process and a generative process, where the diffusion process gradually adds noise at each time step, while the generative process learns how to recover data distribution from the noise distribution using a neural network.

**Figure 1:**
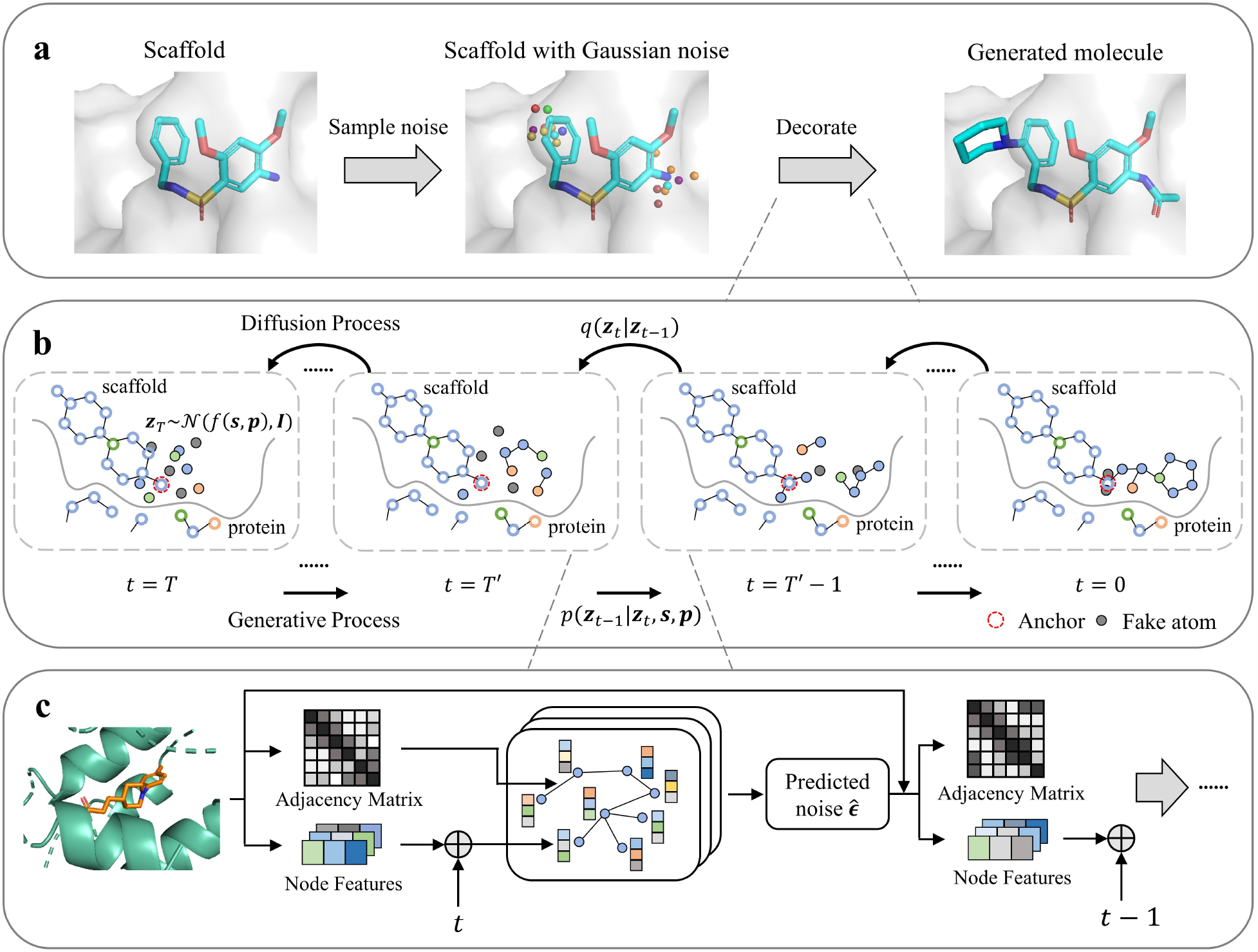
Overview of DiffDec. (a) Flowchart of structure-aware scaffold decoration using DiffDec. (b) Implementation details of DiffDec. The diffusion process iteratively adds Gaussian noise to the data, while the generative process gradually denoises the noise distribution under the condition of scaffold and protein pocket to recover the ground truth R-groups. (c) A detailed denoising procedure from time step *t* to *t −* 1 using EGNN.

### Diffusion process

Following other recently developed diffusion-based models for molecule generation,^37,42^ we iteratively add noise to the data points of R-groups ***x*** using a Gaussian distribution 𝒩 at each time step, but leave the corresponding context ***c*** unchanged throughout the diffusion process. We depict the latent noised representation using ***z***_*t*_ for *t* = 0, …, *T* . Given the previous state ***z***_*t−*1_, the conditional distribution of the data state ***z***_*t*_ at time step *t* is formulated by the multivariate normal distribution:

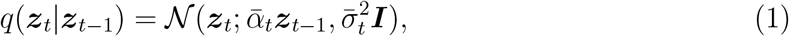

where 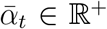 decides how much of the original signal to keep and 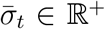 controls how much noise to add. Since the full diffusion process is Markov, the entire noising process can be written as:

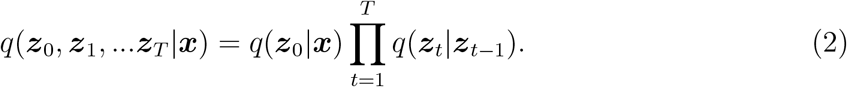

For the distribution *q* is normal and independent at each time step, we can obtain the derivation for the distribution of ***z***_*t*_ given ***x*** as follows:

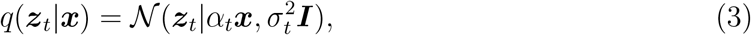

where 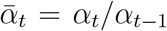 and 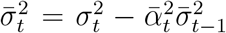. Following Kingma et al.,^43^ we can obtain the signal-to-noise ratio 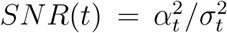. In general, *α*_*t*_ usually follows either a learned or pre-defined schedule from *α*_0_ *≈* 1 towards *α*_*T*_ *≈* 0.^43^ And we can also follow a variance-preserving diffusion process, where 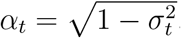.^44^ This closed-form formula indicates that we can achieve the noisy data distribution at any time step without iterative calculations.

With Bayes theorem, the posterior of the denoising transitions from time step *t* to *t –* 1 can be computed in closed-form conditioned on ***x***:

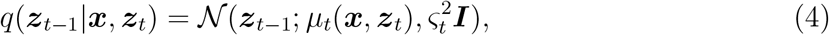

where 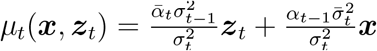 and 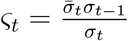.

For the continuous atom coordinates, a non-zero distribution invariant to the translation can not be obtained.^18^ However, we can use the data distributions on the linear subspace where the center of mass (COM) of the system is zero, and this strategy has been used consistently in diffusion.^45,46^ In actuality, this is accomplished by translating the center of mass of the system *f* (***c***) before diffusion process or generative process. Then, the noise can be added from *𝒩* (0, ***I***) instead of *𝒩* (*f* (***c***), ***I***). The choice of *f* is highly related to the problem being solved and prior knowledge. In our problem, we define the atoms connected to the R-groups in the scaffold as anchors. Therefore, if the information of anchors is available, *f* (***c***) can be defined as the coordinates of anchors, and the diffusion processes of multiple anchors are separate and independent of each other; if not, *f* (***c***) can be the center of mass of the scaffold.

For the discrete atomic features, there are also quantities of categorical diffusion models, where we prefer the strategy which places the discrete features on a continuous space. ^36,46^ We encode the variables with one-hot representation and directly add Gaussian noise to it.

To incorporate R-group size prediction into the diffusion model in an end-to-end fashion, we introduce the fake atom mechanism. In general, the size of the R-group does not exceed 10. R-groups with a size smaller than 10 are filled with fake atoms to reach the size of 10 before the diffusion process. The coordinates of these fake atoms are set to *f* (***c***), and an additional dimension of atomic type is used to indicate them. The subsequent noising steps remain the same as described previously. By using fixed-size noisy R-groups, denoising can be performed and the sizes of the R-groups can be determined during the generative process, eliminating the need for an additional prediction module.

### Generative process

The generative process is the reverse of the diffusion process. It will gradually denoise the Gaussian noise obtained from sampling under the fixed context of scaffold and pocket to recover the ground truth R-groups. The generative transition distribution is defined as:

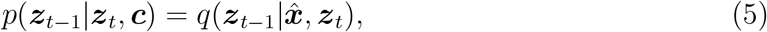

where 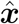 is an approximation of the data point. We use a neural network *φ* to predict the Gaussian noise 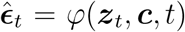 added from time step *t −* 1 to *t* to estimate 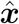.^41^ Therefore, we can obtain:

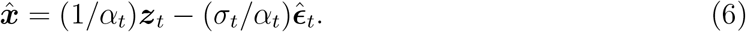

The neural network *φ* mentioned above is implemented as an adjusted E(n)-equivariant graph neural network (EGNN).^47^ It uses the noise-added R-groups ***z***_*t*_, the context ***c*** and the time step *t* as input, and outputs the Gaussian noise that added from time step *t –* 1 to *t*, including coordinates and features 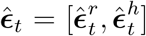. Since the output noises must be on a subspace with a zero center of gravity, the coordinate noises 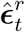should be projected down by subtracting the initial coordinates:^46^

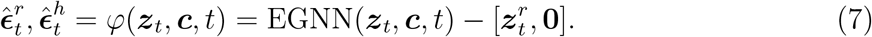

EGNN is made up of *L* layers of Equivariant Graph Convolutional Layers (EGCL), ***r***^*l*+1^, ***h***^*l*+1^ = EGCL[***r***^*l*^, ***h***^*l*^]. Specifically, at the *l*th layer, the atom coordinates ***r*** and features ***h*** are updated alternately as follows:

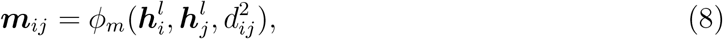

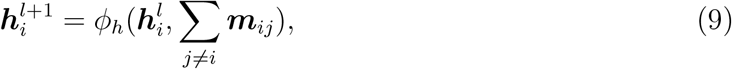

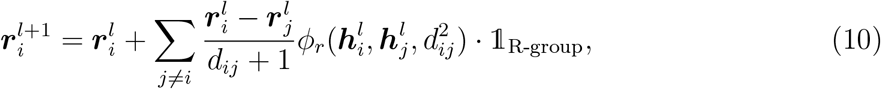

where 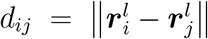 is the Euclidean distance between nodes *i* and *j, ϕ*_*m*_, *ϕ*_*h*_, *ϕ*_*r*_ are learnable components parameterized by fully connected neural networks and 𝟙_R-group_ is the R-group mask (the R-groups are 1 while the contexts are 0) since we want to keep the context coordinates fixed throughout the EGCL layers. The equivariance of EGCL can be inferred from its inputs. For the update of atom features, *ϕ*_*m*_ and *ϕ*_*h*_ is only related to node features ***h*** and distance *d*_*ij*_ between nodes *i* and *j*, which are E(3)-invariant. For the update of atom coordinates, it is related to the difference between ***r***_*i*_ and ***r***_*j*_, which preserves E(3)-equivariance. Therefore, EGNN, which is composed of EGCL, is also E(3)-equivariant.

For graph construction, we assign edges to atom pairs that are closer than 4Å in 3D space. This ensures that the resulting graph is not too dense, reducing computational costs while allowing for messages to be passed effectively.

After passing messages through the EGCL layers, we can get new coordinates 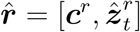 that aggregate information about neighbours as well as features 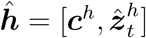. We focus solely on the generated R-groups and disregard the contextual information in the graph. Therefore, the output of the EGNN specifically captures the information pertaining to the R-groups 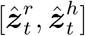.

During the generative process, the initial noise can be sampled from *𝒩* (0, ***I***) instead of *𝒩* (*f* (***c***), ***I***) due to the introduction of *f* (***c***) in the diffusion process. Moreover, during the multi R-groups generative process, instead of treating each anchor’s generation task sequentially, we incorporate the R-groups of the remaining anchors obtained from the previous time step into the context during the message-passing process. This allows us to effectively denoise one anchor’s R-group by leveraging the collective information from all anchors.

### Model training

The model can be trained by optimizing the variational lower bound (VLB):^43,46^

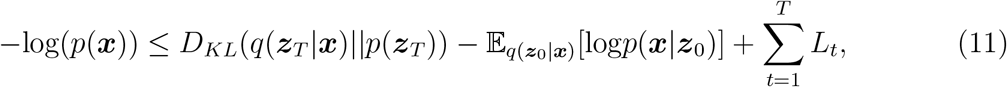

where the first term is the prior loss, the second term is the reconstruction loss and the third term is diffusion loss

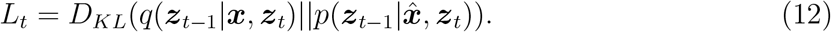

It should be noticed that the prior loss without learnable parameters is always close to zero while the reconstruction loss must be estimated. ^46^ Thus, we instead minimize the simplified training objective, i.e. the mean squared error between true noise and predicted noise:^36,41,43^

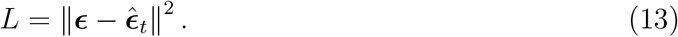

DiffDec was implemented in Python (version 3.10). The models were trained on one GPU (Quadro RTX 5000) for 1,000 steps and saved checkpoints per 100 steps. When training the model, batch size, learning rate, EGCL layers, hidden states, and aggregation method were set to 16, 0.0002, 6, 128, and summation, respectively. All generated candidate molecules were canonicalized using RDkit^48^ and compared to the ground truth molecules. Ablation studies in the Results section are based on three models trained with different pocket information and six models trained with different R-group size information. All these models were trained separately.

### Evaluation metrics

We assessed the generated molecules with a range of metrics:

- **Validity** refers to the percentage of generated molecules which preserve the original scaffolds and can be parsed by RDkit.
- **Uniqueness** is the proportion of unique compounds in generated ones.
- **Recovery** counts the percentage of generated molecules that recover the reference molecules in the test set.
- **Similarity** means the average Tanimoto similarity^49^ between the generated molecules and reference molecules in the test set.
- **Vina score** estimates the binding affinity between the generated molecules and the target protein. The Vina score (kcal/mol) is calculated by QVina.^50^
- **High affinity** is the percentage of test cases where the Vina scores of the generated molecules are higher than or equal to that of the reference compound.

### Baselines

We compared our approaches with the following baselines:

- LibINVENT.^22^ LibINVENT utilized SMILES strings to encode molecules. It consisted of an encoder-decoder architecture where both the encoder and decoder were recurrent neural networks. It is a tool capable of proposing chemical libraries of compounds with the same scaffolds while maximizing desirable properties using reinforcement learning.
- Pocket2Mol.^32^ Pocket2Mol employed an auto-regressive sampling scheme. It is a GNN-based method that generates 3D molecules atom-wise under the context of the pocket. This method has been successfully employed for the structure-based *de novo* drug design.
- FLAG.^33^ FLAG is a GNN-based model that generates 3D molecules with valid and realistic substructures fragment-by-fragment based on the structure information of proteins. This approach is also applicable to structure-based *de novo* drug design.

Notably, these algorithms were not designed for 3D structure-aware scaffold decoration. Therefore, we adjusted them to our tasks for fair comparison. The experiment setup is shown in Supporting Information Text S1.

## Results

### DiffDec achieved state-of-the-art performance

We compared our method with three deep generative models on two newly constructed reaction-based scaffold decoration datasets, i.e. the single R-group and multi R-groups datasets. For each pair of scaffold and pocket in the test sets (43 pairs on single R-group dataset and 102 pairs on multi R-groups dataset), we generated 100 molecules for evaluation to avoid random fluctuation.

On the single R-group decoration task, our method achieved the best performances on metrics validity, recovery, and similarity (Table 1). Specifically, our model attained a validity of 98.00%, while LibINVENT, Pocket2Mol, and FLAG maintained 87.35%, 51.14%, and 87.95%, respectively, indicating that our model could preserve the scaffolds better or generate fewer R-groups in conflict with the scaffolds. The significantly lower performance of Pocket2Mol could be attributed to its atom-wise method in an auto-regressive manner to generate atoms and bonds. In terms of the recovery rate, our model exhibited the highest recovery of 69.67%, which was significantly higher than that of LibINVENT (37.33%), Pocket2Mol (4%), and FLAG (0%). These suggested that our method could effectively recover the reference R-groups, which were regarded as realistic decorations for the scaffolds. The worse performance of Pocket2Mol and FLAG maybe due to their auto-regressive generation scheme, which would lead to error accumulation and difficulties in recovery of the reference R-groups at each step. In addition, DiffDec reached a uniqueness of 48.54%, indicating that DiffDec could generate a diverse set of R-groups, not just restricted to the references. For the similarity of generated molecules to the references, DiffDec achieved an average value of 0.86, higher than 0.80, 0.69, and 0.70 by LibINVENT, Pocket2Mol, and FLAG, respectively.

**Table 1:** Performance comparison of four methods on single R-group decoration task.

| Method | Validity<br>(%, ↑) | Uniqueness<br>(%, ↑) | Recovery<br>(%, ↑) | Similarity<br>(↑) | Vina Score<br>(kcal/mol, ↓) | High Affinity<br>(%, ↑) |
| --- | --- | --- | --- | --- | --- | --- |
| Scaffolds | - | - | - | - | -7.73 | - |
| Reference molecules | - | - | - | - | -8.47 | - |
| LibINVENT | 87.35 | 41.05 | 37.33 | 0.80 | -8.10 | 43.76 |
| Pocket2Mol<sup>a</sup> | 51.14 | 44.27 | 4.00 | 0.69 | -8.11 | <u>44.18</u> |
| FLAG<sup>a</sup> | 87.95 | **65.30** | 0.00 | 0.70 | -7.62 | 34.53 |
| DiffDec (w/o anchors) | <u>93.05</u> | <u>55.33</u> | <u>60.67</u> | <u>0.83</u> | <u>-8.16</u> | 42.11 |
| DiffDec | **98.00** | 48.54 | **69.67** | **0.86** | **-8.25** | **55.06** |

<sup>a</sup> Pocket2Mol and FLAG were adjusted to the structure-aware scaffold decoration task.

To evaluate the binding performance of generated molecules in the target proteins, we docked the compounds to the target protein pockets and computed the corresponding binding affinities, i.e. Vina scores. As expected, our method showed the best performance, with an average docking score of -8.25 kcal/mol and about 55.06% of generated molecules exhibited higher affinity than the reference ligand, compared with those of LibINVENT (-8.10 and 43.76%), Pocket2Mol (-8.11 and 44.18%), and FLAG (-7.62 and 34.53%). Then, we compared the binding performance of these methods in different cases in detail (Figure 2a). In most cases, the average docking score differences of DiffDec were significantly larger than the other three baselines. The molecules generated from DiffDec exhibited the best binding affinity in about 52% cases, while the ones from LibINVENT, Pocket2Mol, and FLAG were best for 8%, 28%, and 12% of all cases.

**Figure 2:**
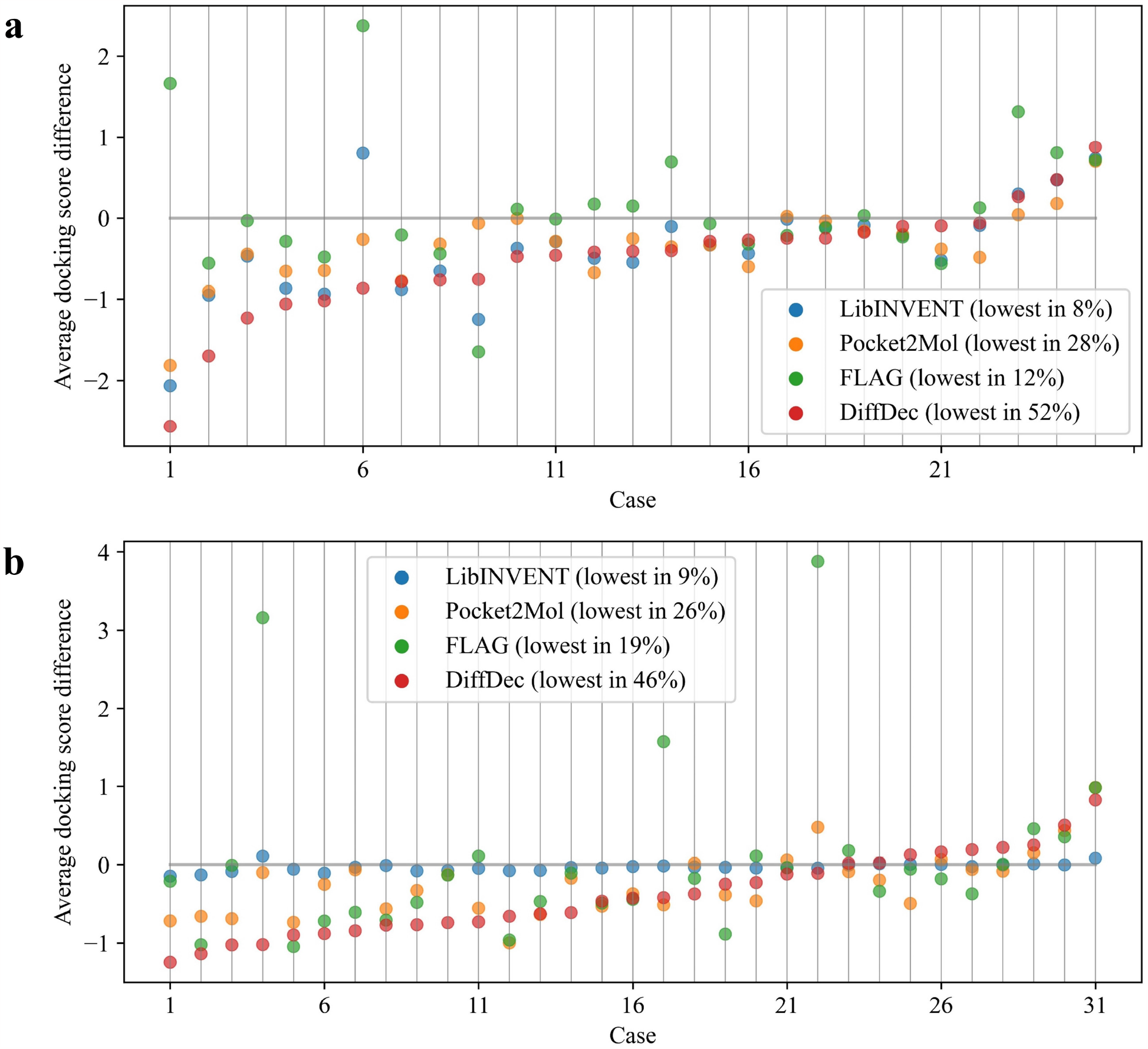
Average Vina score differences between generated molecules and scaffolds across (a) single R-group test cases and (b) multi R-groups test cases.

On the multi R-groups decoration task, the trend was roughly the same as that of the single R-group task: our DiffDec model still achieved the best performance (Table 2) with validity of 97.13%, recovery of 45.34%, similarity of 0.80, Vina score of -8.19 kcal/mol and high affinity of 40.50%. Among these methods, LibINVENT exhibited a low validity and uniqueness of 8.62% and 6.93%. This is because LibINVENT was prone to generating invalid sequences that did not conform to the SMILES syntax rules. For the binding performance comparison of these models, Figure 2b showed that the molecules generated by DiffDec exhibited the best binding affinity in large amounts of the targets (46% vs 9%, 26%, 19% for LibINVENT, Pocket2Mol, FLAG respectively). These results implied that DiffDec still exhibited strong scaffold decoration capability in the more challenging multi R-groups decoration task.

**Table 2:**
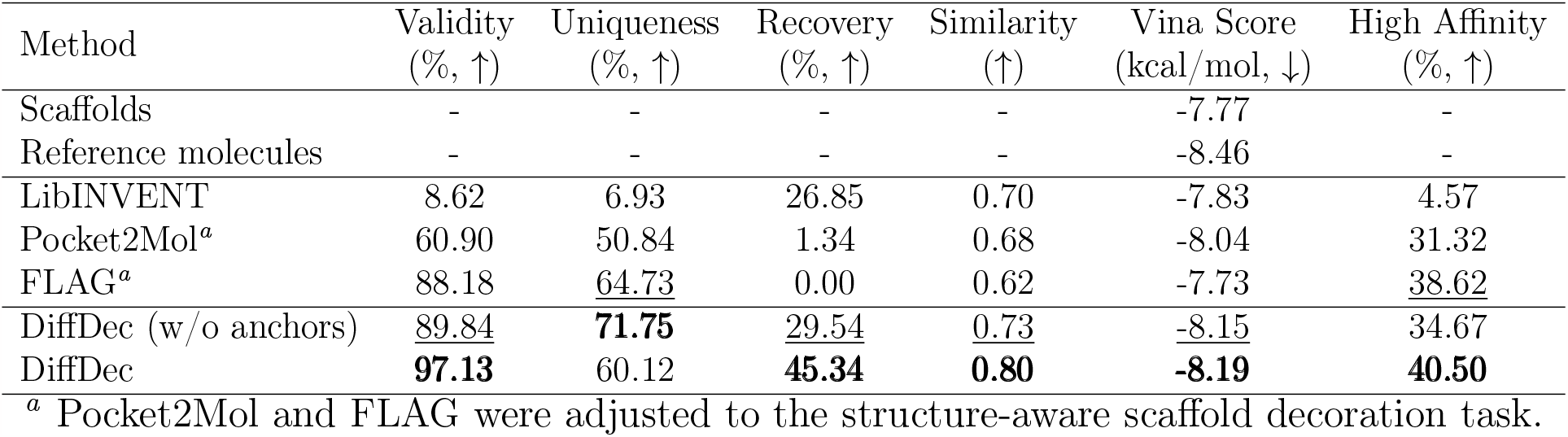
Performance comparison of four methods on multi R-groups decoration task.

We further visualized the distributions of R-groups on the single and multi R-groups decoration tasks. As shown in Figure 3a-d, most of the R-groups generated by LibINVENT and FLAG appeared in the training set, while Pocket2Mol generated a large proportion of compounds outside of the training sets due to its sequential atom-by-atom generation strategy that would lead to unrealistic unseen substructures. On the contrary, DiffDec achieved a balanced result to keep the distribution of training data and to generate novel R-groups outside the training space. On the multi R-groups decoration task, all methods showed similar distributions except LibINVENT which generated quite a small number of R-groups (Figure 3e) due to the low validity (8.62%). These results demonstrated that DiffDec could primely recover the R-groups in the training set and also generate novel molecules.

**Figure 3:**
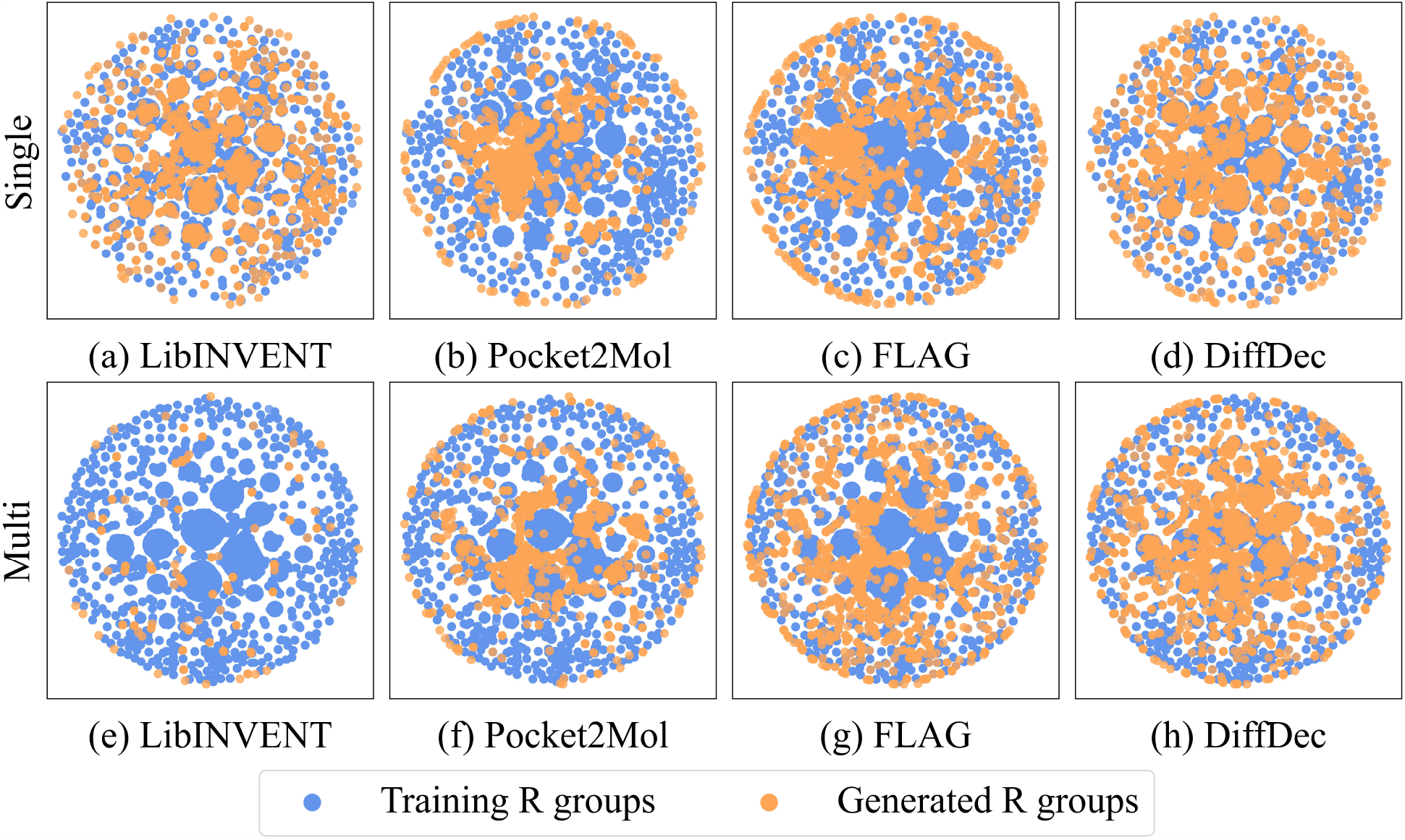
The t-SNE plots of training R-groups and generated R-groups from LibINVENT, Pocket2Mol, FLAG, and DiffDec on single and multi R-groups decoration tasks.

#### DiffDec could generate R-groups well without provided anchors

In the actual scenarios of lead optimization, the specific location of anchors are usually not known, so we investigated the R-groups generation performance of DiffDec without provided anchors (w/o anchors). When not providing anchors information, on the single R-group decoration task, DiffDec (w/o anchors) achieved validity, recovery, and similarity of 93.05%, 60.67%, and 0.83, which were only 4.95%, 9.00%, and 0.03 lower than DiffDec (98.00%, 69.67%, and 0.86) (Table 1). For binding affinity of generated molecules to the target pocket, DiffDec (w/o anchors) exhibited the average Vina score and high affinity percentage of -8.16 kcal/mol and 42.11%, which were comparable to those of DiffDec (-8.25 and 55.06%). The reason for these declines is that DiffDec without anchors information needed to consider both the generation of the R-group and the appropriate anchor to attach it to, which resulted in more challenging generation. On the multi R-groups decoration task, DiffDec (w/o anchors) achieved validity of 89.84%, recovery of 29.54%, similarity of 0.73, Vina score of -8.15 kcal/mol, and high affinity of 34.67%, which decreased by 7.29%, 15.80%, 0.07, -0.04, and 5.83% compared to DiffDec (97.13%, 45.34%, 0.80, -8.19, and 40.50%) (Table 2). Even so, DiffDec (w/o anchor) outperformed the other three baseline models with anchor information on most metrics. Therefore, even if not providing anchors, our method could automatically identify anchors and generate R-groups with acceptable quality and comparable performance.

#### The importance of pocket information

To illustrate DiffDec’s awareness of pocket information, we retrained DiffDec and evaluated the performances under two conditions: with pocket backbone information and without any pocket information. We calculated the number of clashes between the generated R-groups and surrounding pockets for comparison. The clash is defined as if the distance between two atoms is less than the sum of their van der Waals radius. As shown in Figure 4a, the number of clashes generated by DiffDec with full-atomic pocket representation (with an average of 8 clashes per molecule) was almost identical to that of the reference complex in the test set (with an average of 7 clashes per molecule). When the protein pocket was represented as the backbone or removed, the average number of clashes was increased to 15 or 18 respectively. Then, we performed R-groups generation for ABL2/ARG kinase (4xli)^51^ as an example by DiffDec without pockets and with pockets as the context for comparison (Figure 4b). When using DiffDec without pocket information, the generated R-groups collided with the surrounding amino acid residues in the pocket (the red dashed box). While in DiffDec with full-atomic pocket information, the R-groups were generated according to the condition of the surrounding pockets and orientated to an appropriate position to make fewer clashes with the pocket.

**Figure 4:**
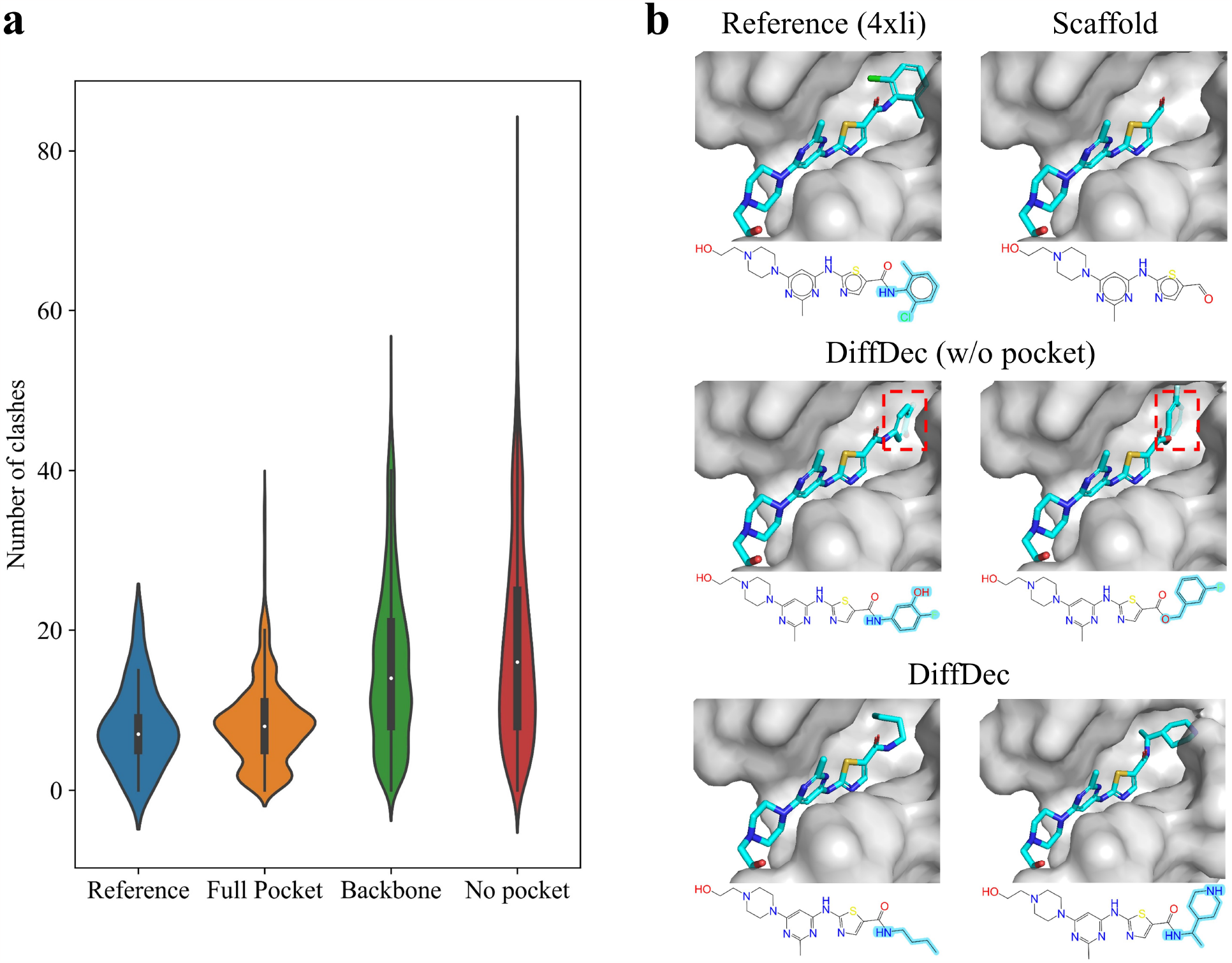
Results of the ablation study of pocket information. (a) Distribution of numbers of clashes in reference test set and molecules sampled from DiffDec models with different pocket representations. (b) R-groups generated by DiffDec without pockets and with pockets as context (PDB id is 4xli).

#### The importance of the fake atom module

To illustrate the importance of the fake atom module in DiffDec, we removed the module from DiffDec and utilized a trained graph neural network to predict the size of R-groups (size GNN). By comparison, we showed the results of DiffDec using the R-group sizes in ground truth (given size). As shown in Table 3, DiffDec with size GNN achieved recovery of 54.67%, Vina score of -8.15 kcal/mol, and high affinity ratio of 49.89%. These values were significantly lower than those of DiffDec (69.67%, -8.25 kcal/mol, and 55.06%) and DiffDec (given size) (79.33%, -8.29 kcal/mol, and 60.97%). In addition, the DiffDec (w/o anchors) model also exhibited better recovery ratio, Vina score, and high affinity ratio than DiffDec (w/o anchors, size GNN). These results suggested that the fake atom module could predict a better R-group size than GNN to accommodate the surrounding pocket and benefit the subsequent generation.

**Table 3:** Ablation study of the fake atom module on single R-group decoration task.

| Method | Validity<br>(%, ↑) | Uniqueness<br>(%, ↑) | Recovery<br>(%, ↑) | Similarity<br>(↑) | Vina Score<br>(kcal/mol, ↓) | High Affinity<br>(%, ↑) |
| --- | --- | --- | --- | --- | --- | --- |
| DiffDec (w/o anchors, size GNN) | **95.51** | 55.23 | 51.67 | 0.82 | -8.09 | 38.85 |
| DiffDec (w/o anchors) | 93.05 | **55.33** | <u>60.67</u> | <u>0.83</u> | <u>-8.16</u> | <u>42.11</u> |
| DiffDec (w/o anchors, given size) | 95.16 | 38.33 | **62.00** | **0.85** | **-8.18** | **46.57** |
| DiffDec (size GNN) | 97.87 | 48.27 | 54.67 | 0.85 | -8.15 | 49.89 |
| DiffDec | <u>98.00</u> | **48.54** | <u>69.67</u> | <u>0.86</u> | <u>-8.25</u> | <u>55.06</u> |
| DiffDec (given size) | **98.07** | 41.26 | **79.33** | **0.88** | **-8.29** | **60.97** |

#### Case study

We provided several case studies for DiffDec on single and multi R-groups generation and showed examples of generated molecules with higher binding affinities (lower Vina score) than the reference compounds (Figure 5, S2-S3). As shown in Figure 5, S2-S3, DiffDec could generate compounds that bound geometrically complementary to the target pocket, and the generated molecules exhibited significantly improved binding affinities compared to the scaffold and reference ligands. Then we analyzed the binding modes and interactions of molecule 1-1 and 1-2 with surrounding amino acid residues using Maestro of Schrödinger^52^ to interpret the binding affinity improvement of these generated molecules. Compared to the reference molecule, the generated R-groups of molecule 1-1 and 1-2 formed an additional hydrogen bond with Asn220 of the protein, and mol 1-1 formed another additional H-bond interaction with Asn218 (Figure 5). Thus, the binding affinities of molecule 1-1 and 1-2 were higher than those of the reference ligand. When decorating multi R-groups (Figure 5 mol2-1 and mol2-2), DiffDec could also generate molecules with significantly improved affinities and recover the complex trifluoromethyl group in the reference compound. Therefore, DiffDec is capable of generating molecules with improved binding affinities and more favourable binding interactions for lead optimization.

**Figure 5:**
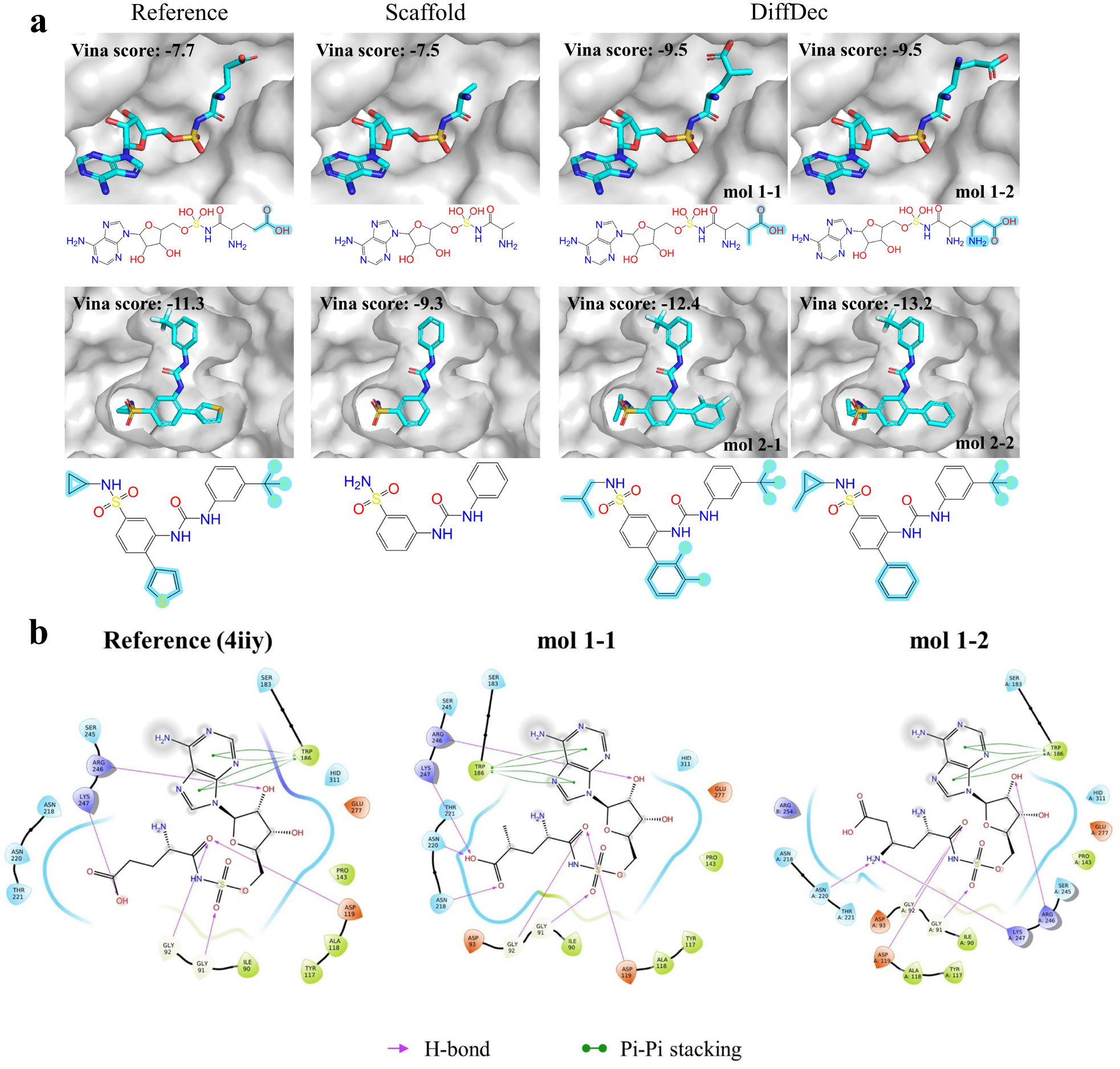
(a) R-groups generated by DiffDec on single and multi R-groups decoration tasks. The two rows represent two different scaffolds and pockets (PDB ids are 4iiy and 4ja8 respectively). (b) Protein-ligand interaction of case 4iiy (PDB id is 4iiy).

## Discussion and conclusions

In this study, we developed an E(3)-equivariant 3D-conditional diffusion model, DiffDec, for molecular scaffold decoration conditioned on 3D protein pockets. The model could generate single and multi R-groups for scaffold decoration inside protein pockets based on two newly constructed benchmark datasets.

Comprehensive evaluations demonstrated that DiffDec achieved state-of-the-art performance on R-groups generation and outperformed the baseline methods on conventional metrics such as validity, recovery, similarity, and binding affinity. The t-SNE projection indicated that our model could not only well recover the R-groups in the training set but also generate a number of novel R-groups out of learning space. In addition, our method could identify the growth anchors and generate R-groups well for scaffolds without provided anchors. Besides, our model conditioned on full-atomic pocket representation generated R-groups with a minimum number of clashes. Our newly introduced fake atom module could predict the suitable R-groups sizes according to the surrounding pocket. Case studies indicated that DiffDec generated decorated molecules which were geometrically complementary to the target pocket and exhibited higher binding affinities and more favourable interactions with the surrounding pocket than the reference compound.

However, there are still some limitations of DiffDec in the current framework. First, the output of DiffDec is a 3D atomic point cloud of the generated R-groups, and the covalent bonds between pairs of atoms are additionally computed by a bond inference algorithm (e.g. OpenBabel^53^). Therefore, the accuracy of OpenBabel will impact the validity and recovery ratio of the generated molecules. Optimization of this bond generation problem could be the direction of our further work. Second, the R-groups generation process in DiffDec requires step-by-step denoising from Gaussian noise, which is computationally intensive and limits the sampling efficiency. This may be solved by combining distributed CPU and GPU computations in a mixed fashion.

In summary, DiffDec provides a flexible and effective method for 3D structure-aware scaffold decoration. We believe that our method will provide a practical way in real scenarios and inspire future scaffold decoration work.

## Acknowledgement

The authors thank Dr. David Daqiang Xu for providing useful suggestions for paper revisions. This work has been supported by the National Key R&D Program of China (2020YFB0204803), the Guangzhou S&T Research Plan (202007030010), the National Natural Science Foundation of China (82003651), Guangdong-Hong Kong Technology Cooperation Funding Scheme (2023A0505010015), and Guangzhou Basic and Applied Basic Research Project (202201011795).

## Supporting Information Available

The supporting Information is available at suppinfo.pdf.

Text S1, experiment setup of the other methods; Table S1, data statistics for the Cross-Docked, single R-group and multi R-groups datasets; Figure S1, distributions of R-groups’ size; and Figures S2, S3, example generations of each task.

