## Supporting Information for "DiffDec: Structure-Aware Scaffold Decoration with an End-to-End Diffusion Model"

### Text S1: Experiment setup of the other methods

LibINVENT[1] is a SMILES-based deep generative model for scaffold decoration, so we only converted the molecular graphs in our datasets into SMILES sequences and inputted them into LibINVENT without the corresponding protein information. Then LibINVENT would generate a batch of decorated molecules.

Pocket2Mol[2] and FLAG[3] are both structure-based de novo drug design methods. For a fair comparison, we adjusted them to our scaffold decoration task and made two major changes. First, in addition to the target pocket, we also input the molecular scaffold which was set in advance and supposed to have been generated. Second, we limited the number of generated R-groups and restricted the generation on the R-groups' attachment points and subsequent atoms or fragments.

Then, we retrained the three models on our single and multi R-groups datasets following the same setup as reported in their studies.

Table S1: Data statistics for the CrossDocked, single R-group and multi R-groups datasets.

| | CrossDocked | Training set (single) | Test set (single) | Training set (multi) | Test set (multi) |
|---|---|---|---|---|---|
| #Molecules | 8472 | 4258 | 25 | 4232 | 31 |
| #Scaffolds | - | 7127 | 43 | 10584 | 102 |
| #R-groups | - | 837 | 35 | 886 | 44 |

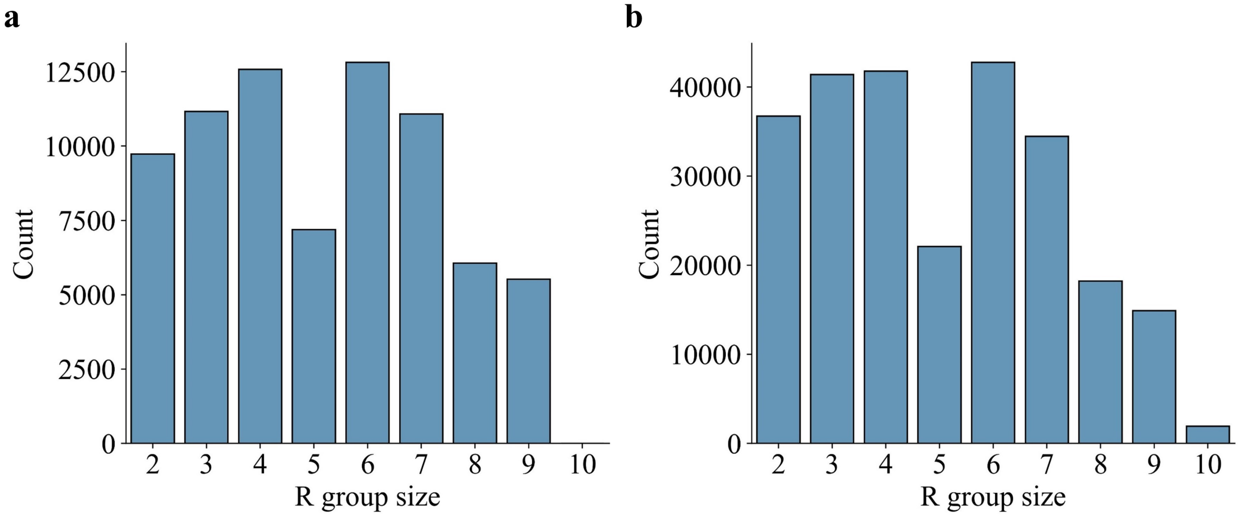


Figure S1: (a) Distribution of size of R-groups on the single R-group decoration dataset. (b) Distribution of size of R-groups on the multi R-groups decoration dataset.

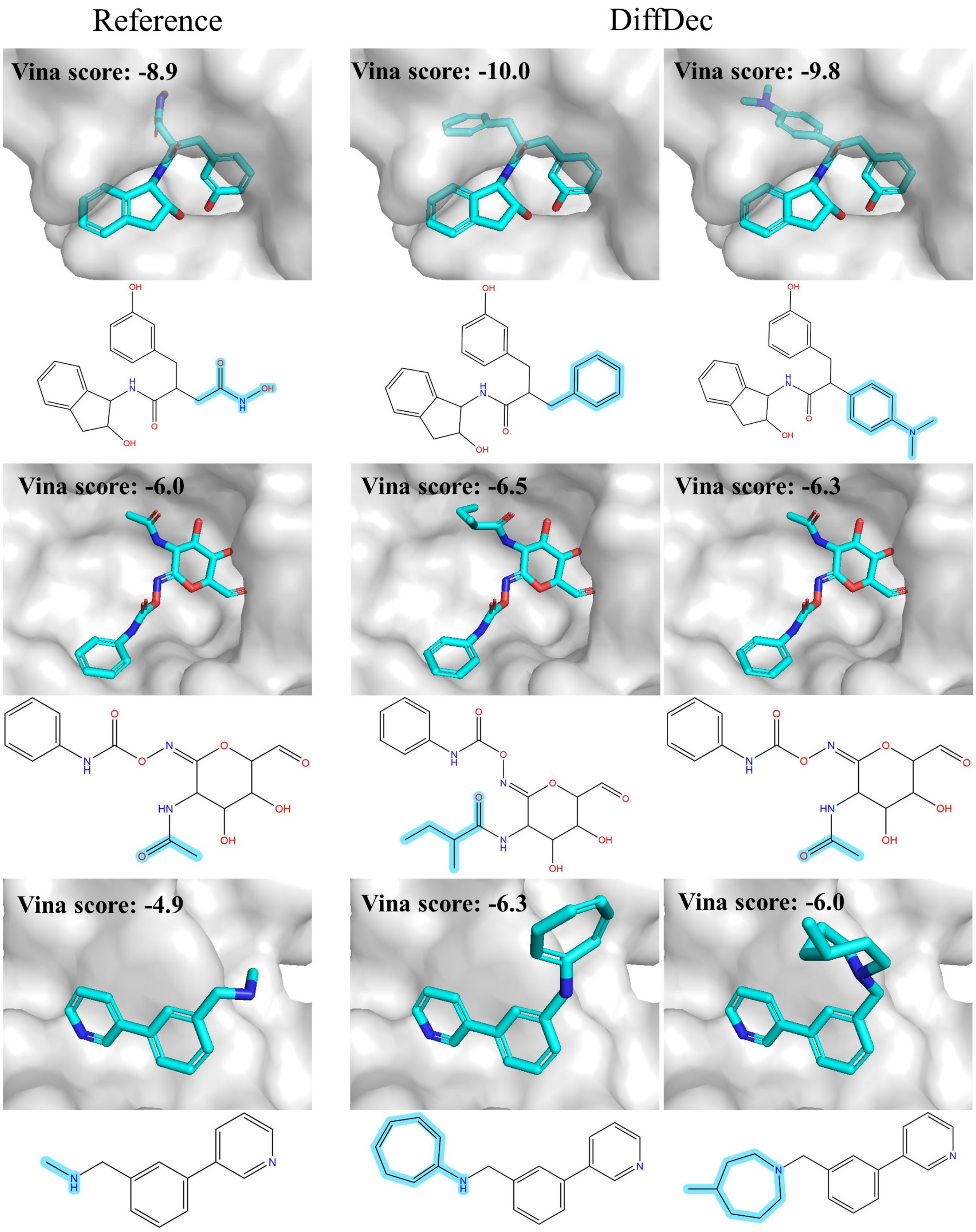


Figure S2: Examples of R-groups generated by DiffDec on the single R-group decoration task (PDB ids are 3hy9, 3gs6 and 4u5s respectively).

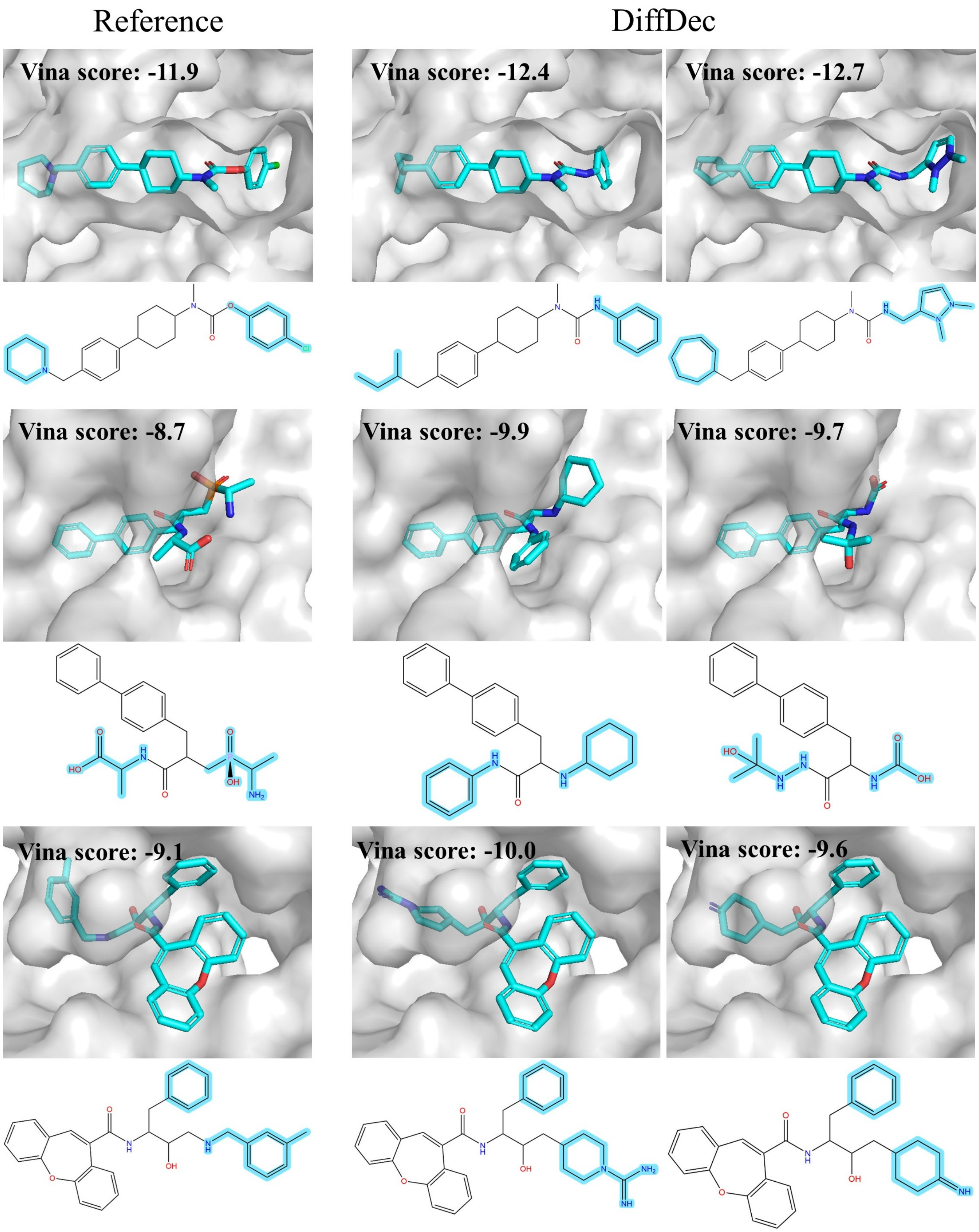


Figure S3: Examples of R-groups generated by DiffDec on the multi R-groups decoration task (PDB ids are 1h36, 1r1h and 4bel respectively).
